## Supplementary Materials for "Pollen Patterns Form from Modulated Phases"

**Supplemental Data**

- Figure S1: Detailed phylogenetic tree with 202 labeled families in spermatophytes. Colored dot represent the character states of the species within each family. Each terminal taxon is labeled with up to four states. The scale bar represents 40.0 million years.
- Table S1: List of papers used to categorize pollen.
- File S1: Glycosyl composition analysis completed at the Complex Carbohydrate Research Center at the University of Georgia for *Passiflora incaranta*.
- File S2: Glycosyl linkages analysis completed at the Complex Carbohydrate Research Center at the University of Georgia for *Passiflora incaranta*.
- File S3: Nexus file used in our analysis. This file includes the morphological data for all spermatophyte families and the phylogenetic tree.

**Figure S1: Phylogenetic Tree with Family Names**

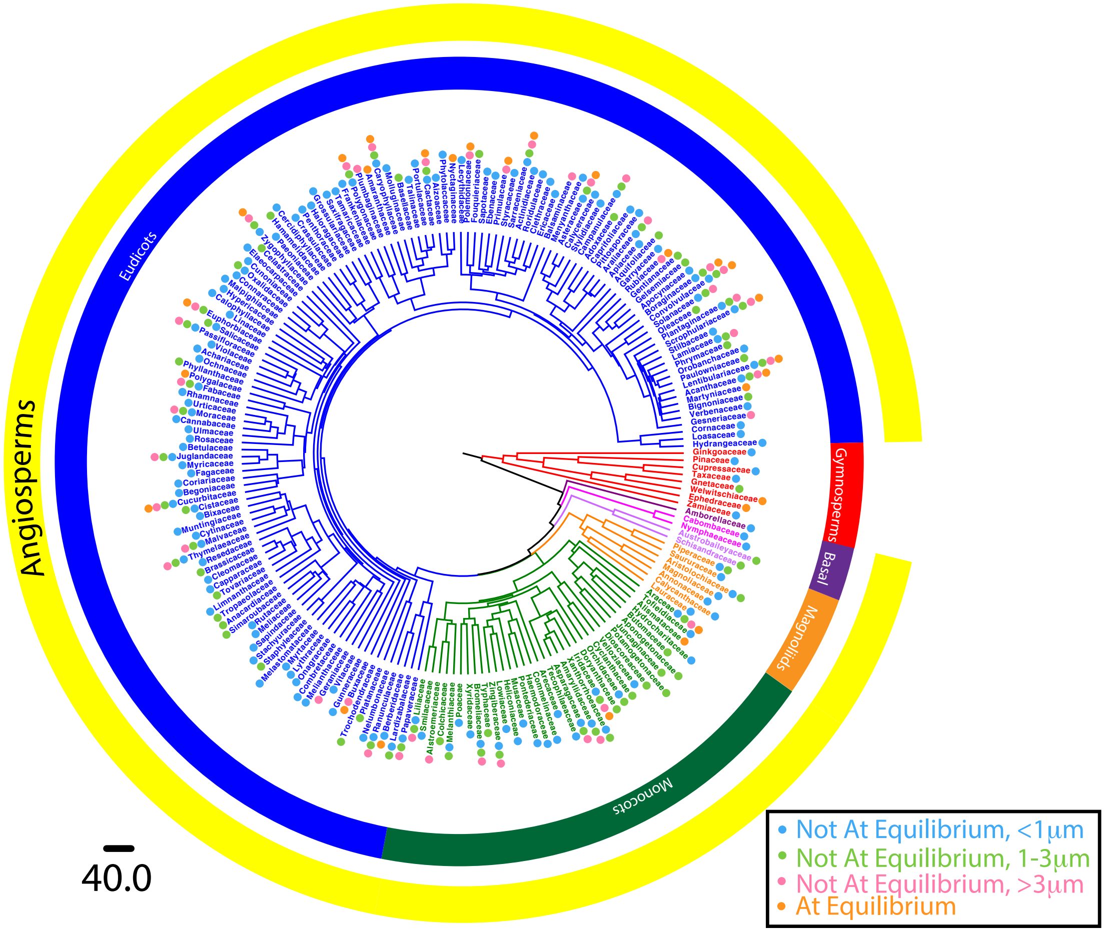


**Supplemental Figure 1: Detailed Angiosperm Phylogenetic Tree with Character States.** Phylogenetic tree of spermatophytes with 202 families at terminal taxa. Colored dot represent the character states of the species within each family. Each terminal taxon is labeled with the family name and with up to four states. The numbered families (27 in total) are those which have species that are in an equilibrium states. The families listed in black have more than one state; families listed in bold (seven of the 27) only have species that are in an equilibrium state. The scale bar represents 40.0 million years.

**Table S1: List of papers used to categorize pollen**

Literature used to categorize pollen species into equilibrium or non-equilibrium states

| Family | References |
|---|---|
| Annonaceae | Gabarayeva 1995 |
| Araceae | Anger & Weber 2006, Weber et al. 1998 |
| Aristolochiaceae | Gonzalez et al 2001, Polevova 2015 |
| Asteraceae | Blackmore & Barnes 1988, Blackmore et al. 2010, Tomb et al. 1974, Takahashi 1989, Dickinson & Potter 1976, Blackmore & Barnes 1987, Horner Jr. & Pearson 1978 |
| Austrobaileyaceae | Zavada 1984 |
| Boraginaceae | Gabarayeva et al. 2011 |
| Brassicaceae | Fitzgerald & Knox 1995 |
| Caryophyllaceae | Heslop-Harrison 1963, Audran & Batcho 1981, Shoup et al 1980 |
| Convolvulaceae | Echlin et al. 1966 |
| Ephedraceae | Doores et al. 2007 |
| Gnetaceae | Yao et al, 2004 |
| Heliconiaceae | Stone 1987 |
| Hydrocharitaceae | Takahashi 1994 |
| Lauraceae | Stone 1987, Rowley & Vasanthy 1993 |
| Liliaceae | Sheldon & Dickinson 1986, Heslop-Harrison 1968 |
| Malvaceae | Takahashi & Kouchi 1988 |
| Nyctaginaceae | Takahashi & Skvarla 1991 |
| Nymphaeaceae | Takahashi 1992 |
| Papaveraceae | Romero et al 2003 |
| Welwitschiaceae | Doores t al. 2007 |

**File S1: Glycosyl composition analysis completed at the Complex Carbohydrate Research Center at the University of Georgia for *Passiflora incaranta*.**

**Date:** 2/28/17

**Investigator:** **Asja Radja**
University of Pennsalvania
209 S 33rd Street
Philadelphia, PA 19104

**Subject:** Glycosyl composition analysis.

**Sample** **Sample #1**

**CCRC Code:** **AR021617**

**Analyst:** Ian Black

**Methods:**

**Should any of these data be used in a publication, please include the following statement in the acknowledgment: This work was supported by the Chemical Sciences, Geosciences and Biosciences Division, Office of Basic Energy Sciences, U.S. Department of Energy grant (DE-SC0015662) to Parastoo Azadi " at the Complex Carbohydrate Research Center.**

Glycosyl composition

Glycosyl composition analysis was performed by combined gas chromatography/mass spectrometry (GC/MS) of the per-*O*-trimethylsilyl (TMS) derivatives of the monosaccharide methyl glycosides produced from the sample by acidic methanolysis as described previously by Santander *et al*. (2013) *Microbiology* **159**:1471.

Briefly, the sample (300 ug) was heated with methanolic HCl in a sealed screw-top glass test tube for 17 h at 80 °C. After cooling and removal of the solvent under a stream of nitrogen, the sample was treated with a mixture of methanol, pyridine, and acetic anhydride for 30 min. The solvents were evaporated, and the sample was derivatized with Tri-Sil® (Pierce) at 80 °C for 30 min. GC/MS analysis of the TMS methyl glycosides was performed on an Agilent 7890A GC interfaced to a 5975C MSD, using an Supelco Equity-1 fused silica capillary column (30 m × 0.25 mm I

### Glycosyl Composition Analysis

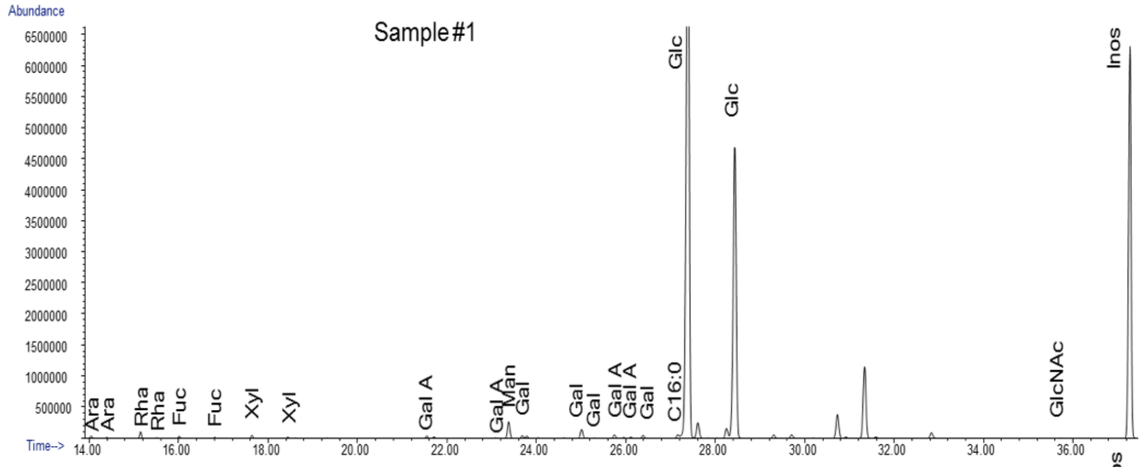


The GC chromatograms of the compositional analysis.

| Sample | Glycosyl residue | Mass ([illegible] g) | Mol % |
|---|---|---|---|
| Sample #1 | Ribose (Rib) | n.d. | - |
| | Arabinose (Ara) | 0.2 | 0.2 |
| | Rhamnose (Rha) | 0.7 | 0.7 |
| | Fucose (Fuc) | 0.2 | 0.2 |
| | Xylose (Xyl) | 0.4 | 0.4 |
| | Glucuronic Acid (GlcA) | n.d. | - |
| | Galacturonic acid (GalA) | 0.8 | 0.6 |
| | Mannose (Man) | 1.5 | 1.3 |
| | Galactose (Gal) | 1.4 | 1.2 |
| | Glucose (Glc) | 106.9 | 95.2 |
| | N-Acetyl Galactosamine (GalNAc) | n.d. | - |
| | N-Acetyl Glucosamine (GlcNAc) | 0.1 | 0.1 |
| | N-Acetyl Manosamine (ManNAc) | n.d. | - |
| | [illegible] | 112.2 | |

The estimated amounts and mole percentage of each detected monosaccharide in the sample.

**File S2: Glycosyl linkages analysis completed at the Complex Carbohydrate Research Center at the University of Georgia for *Passiflora incaranta*.**

**Date:** 3/6/17

**Investigator:** **Asja Radja**
University of Pennsylvania
209 S 33rd Street
Philadelphia, PA 19104

**Subject:** Glycosyl linkage analysis.

**Sample** **Sample #1**

**CCRC Code:** **AR021617**

**Analyst:** Ian Black

**Methods:**

**Should any of these data be used in a publication, please include the following statement in the acknowledgment: This work was supported by the Chemical Sciences, Geosciences and Biosciences Division, Office of Basic Energy Sciences, U.S. Department of Energy grant (DE-SC0015662) to Parastoo Azadi " at the Complex Carbohydrate Research Center.**

Glycosyl linkage analysis

The glycosyl linkage analysis was performed at the Complex Carbohydrate Research Center and was supported by the Chemical Sciences, Geosciences and Biosciences Division, Office of Basic Energy Sciences, U.S. Department of Energy grant (DE-SC0015662) to Parastoo Azadi.

#### Glycosyl Linkage Analysis

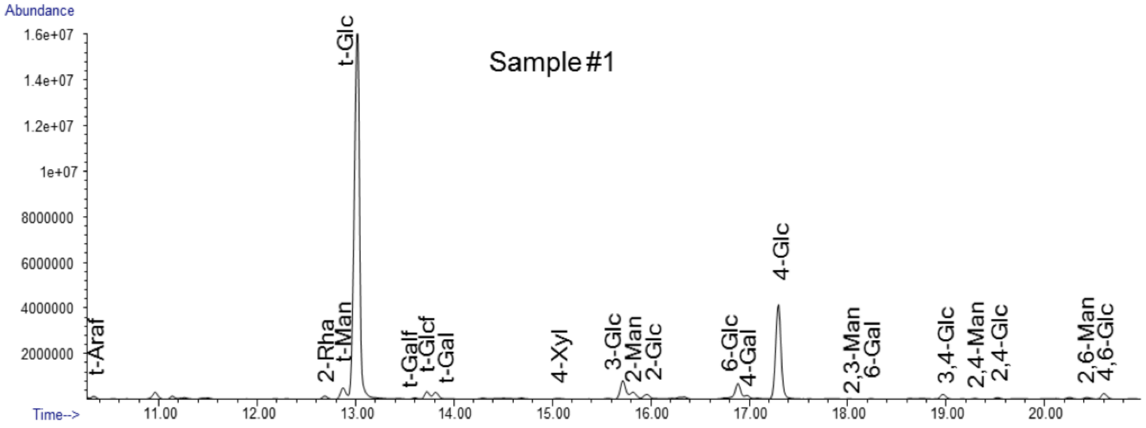


The GC chromatograms of the linkage analysis.

The relative percentage of each detected monosaccharide linkage in the sample.

| Peak | #1 area % |
|---|---|
| Terminal Arabinofuranosyl residue (t-Araf) | 0.4 |
| 2-linked Rhamnopyranosyl residue (2-Rha) | 0.5 |
| Terminal Mannopyranosyl residue (t-Man) | 2.1 |
| Terminal Glucopyranosyl residue (t-Glc) | 66.5 |
| Terminal Galactofuranosyl residue (t-Galf) | 0.2 |
| Terminal Glucofuranosyl residue (t-Glcf) | 1.2 |
| Terminal Galactopyranosyl residue (t-Gal) | 1.2 |
| 4-linked Xylopyranosyl residue (4-Xyl) | 0.1 |
| 3-linked Glucopyranosyl residue (3-Glc) | 3.2 |
| 2-linked Mannopyranosyl residue (2-Man) | 1.4 |
| 2-linked Glucopyranosyl residue (2-Glc) | 1.0 |
| 3-linked Galactopyranosyl residue (3-Gal) | - |
| 6-linked Glucopyranosyl residue (6-Glc) | 2.3 |
| 4-linked Galactopyranosyl residue (4-Gal) | 0.5 |
| 4-linked Glucopyranosyl residue (4-Glc) | 16.8 |
| 2,3-linked Mannopyranosyl residue (2,3-Man) | 0.1 |
| 6-linked Galactopyranosyl residue (6-Gal) | 0.1 |
| 3,4-linked Glucopyranosyl residue (3,4-Glc) | 0.8 |
| 2,4-linked Mannopyranosyl residue (2,4-Man) | 0.1 |
| 2,4-linked Glucopyranosyl residue (2,4-Glc) | 0.2 |
| 2,6-linked Mannopyranosyl residue (2,6-Man) | 0.3 |
| 4,6-linked Glucopyranosyl residue (4,6-Glc) | 1.0 |

**File S3: Nexus File**

```
#NEXUS
begin taxa;
	dimensions ntax=195;
	taxlabels
	10[&!name="gnetaceae",!color=#ff0000]
	101[&!name="cactaceae",!color=#0000ff]
	103[&!name="cornaceae",!color=#0000ff]
	104[&!name="loasaceae",!color=#0000ff]
	105[&!name="hydrangeaceae",!color=#0000ff]
	108[&!name="balsaminaceae",!color=#0000ff]
	11[&!name="welwitschiaceae",!color=#ff0000]
	111[&!name="lecythidaceae",!color=#0000ff]
	112[&!name="polemoniaceae",!color=#0000ff]
	113[&!name="fouquieriaceae",!color=#0000ff]
	114[&!name="sapotaceae",!color=#0000ff]
	115[&!name="ebenaceae",!color=#0000ff]
	116[&!name="primulaceae",!color=#0000ff]
	12[&!name="ephedraceae",!color=#ff0000]
	120[&!name="styracaceae",!color=#0000ff]
	122[&!name="sarraceniaceae",!color=#0000ff]
	123[&!name="actinidiaceae",!color=#0000ff]
	124[&!name="roridulaceae",!color=#0000ff]
	125[&!name="clethraceae",!color=#0000ff]
	127[&!name="ericaceae",!color=#0000ff]
	13[&!name="amborellaceae",!color=#400080]
	134[&!name="aquifoliaceae",!color=#0000ff]
	137[&!name="campanulaceae",!color=#0000ff]
	138[&!name="menyanthaceae",!color=#0000ff]
	140[&!name="asteraceae",!color=#0000ff]
	141[&!name="calyceraceae",!color=#0000ff]
	142[&!name="stylidiaceae",!color=#0000ff]
	15[&!name="cabombaceae",!color=#ff00ff]
	150[&!name="adoxaceae",!color=#0000ff]
	151[&!name="caprifoliaceae",!color=#0000ff]
	155[&!name="pittosporaceae",!color=#0000ff]
	156[&!name="araliaceae",!color=#0000ff]
	158[&!name="apiaceae",!color=#0000ff]
	16[&!name="nymphaeaceae",!color=#ff00ff]
	161[&!name="garryaceae",!color=#0000ff]
	165[&!name="rubiaceae",!color=#0000ff]
	166[&!name="gentianaceae",!color=#0000ff]
	168[&!name="gelsemiaceae",!color=#0000ff]
	169[&!name="apocynaceae",!color=#0000ff]
	17[&!name="austrobaileyaceae",!color=#cc66ff]
	170[&!name="boraginaceae",!color=#0000ff]
	171[&!name="convolvulaceae",!color=#0000ff]
	172[&!name="solanaceae",!color=#0000ff]
```

177[&!name="oleaceae",!color=#0000ff]
181[&!name="gesneriaceae",!color=#0000ff]
182[&!name="plantaginaceae",!color=#0000ff]
183[&!name="scrophulariaceae",!color=#0000ff]
184[&!name="stilbaceae",!color=#0000ff]
185[&!name="lamiaceae",!color=#0000ff]
186[&!name="phrymaceae",!color=#0000ff]
187[&!name="orobanchaceae",!color=#0000ff]
188[&!name="paulowniaceae",!color=#0000ff]
19[&!name="schisandraceae",!color=#cc66ff]
190[&!name="verbenaceae",!color=#0000ff]
191[&!name="bignoniaceae",!color=#0000ff]
193[&!name="martyniaceae",!color=#0000ff]
196[&!name="lentibulariaceae",!color=#0000ff]
198[&!name="acanthaceae",!color=#0000ff]
2[&!name="zamiaceae",!color=#ff0000]
200[&!name="paeoniaceae",!color=#0000ff]
202[&!name="hamamelidaceae",!color=#0000ff]
203[&!name="cercidiphyllaceae",!color=#0000ff]
206[&!name="grossulariaceae",!color=#0000ff]
207[&!name="saxifragaceae",!color=#0000ff]
208[&!name="crassulaceae",!color=#0000ff]
211[&!name="penthoraceae",!color=#0000ff]
212[&!name="haloragaceae",!color=#0000ff]
213[&!name="vitaceae",!color=#000080]
214[&!name="geraniaceae",!color=#000080]
215[&!name="melianthaceae",!color=#000080]
217[&!name="combretaceae",!color=#000080]
218[&!name="onagraceae",!color=#000080]
219[&!name="lythraceae",!color=#000080]
220[&!name="myrtaceae",!color=#000080]
222[&!name="melastomataceae",!color=#000080]
229[&!name="staphyleaceae",!color=#000080]
231[&!name="stachyuraceae",!color=#000080]
237[&!name="anacardiaceae",!color=#000080]
239[&!name="sapindaceae",!color=#000080]
24[&!name="aristolochiaceae",!color=#ff8000]
240[&!name="meliaceae",!color=#000080]
241[&!name="rutaceae",!color=#000080]
242[&!name="simaroubaceae",!color=#000080]
247[&!name="thymelaeaceae",!color=#000080]
248[&!name="malvaceae",!color=#000080]
249[&!name="cytinaceae",!color=#000080]
250[&!name="muntingiaceae",!color=#000080]
252[&!name="bixaceae",!color=#000080]
253[&!name="cistaceae",!color=#000080]
257[&!name="tropaeolaceae",!color=#000080]
26[&!name="piperaceae",!color=#ff8000]

```
261[&!name="limnanthaceae",!color=#000080]
267[&!name="resedaceae",!color=#000080]
268[&!name="tovariaceae",!color=#000080]
27[&!name="saururaceae",!color=#ff8000]
270[&!name="capparaceae",!color=#000080]
271[&!name="cleomaceae",!color=#000080]
272[&!name="brassicaceae",!color=#000080]
273[&!name="zygophyllaceae",!color=#000080]
276[&!name="fabaceae",!color=#000080]
278[&!name="polygalaceae",!color=#000080]
280[&!name="rosaceae",!color=#000080]
283[&!name="rhamnaceae",!color=#000080]
285[&!name="ulmaceae",!color=#000080]
286[&!name="cannabaceae",!color=#000080]
287[&!name="moraceae",!color=#000080]
288[&!name="urticaceae",!color=#000080]
289[&!name="coriariaceae",!color=#000080]
29[&!name="magnoliaceae",!color=#ff8000]
291[&!name="cucurbitaceae",!color=#000080]
293[&!name="begoniaceae",!color=#000080]
298[&!name="fagaceae",!color=#000080]
299[&!name="myricaceae",!color=#000080]
3[&!name="ginkgoaceae",!color=#ff0000]
300[&!name="juglandaceae",!color=#000080]
303[&!name="betulaceae",!color=#000080]
305[&!name="celastraceae",!color=#000080]
307[&!name="connaraceae",!color=#000080]
308[&!name="oxalidaceae",!color=#000080]
309[&!name="cunoniaceae",!color=#000080]
310[&!name="elaeocarpaceae",!color=#000080]
316[&!name="malpighiaceae",!color=#000080]
318[&!name="linaceae",!color=#000080]
32[&!name="annonaceae",!color=#ff8000]
321[&!name="calophyllaceae",!color=#000080]
322[&!name="hyperiacaceae",!color=#000080]
325[&!name="euphorbiaceae",!color=#000080]
327[&!name="phyllanthaceae",!color=#000080]
34[&!name="calycanthaceae",!color=#ff8000]
340[&!name="ochnaceae",!color=#000080]
342[&!name="achariaceae",!color=#000080]
343[&!name="violaceae",!color=#000080]
344[&!name="passifloraceae",!color=#000080]
347[&!name="salicaceae",!color=#000080]
352[&!name="dioscoreaceae",!color=#008040]
354[&!name="velloziaceae",!color=#008040]
357[&!name="cyclanthaceae",!color=#008040]
358[&!name="melanthiaceae",!color=#008040]
360[&!name="colchicaceae",!color=#008040]
```

```
361[&!name="alstroemeriaceae",!color=#008040]
364[&!name="smilacaceae",!color=#008040]
365[&!name="liliaceae",!color=#008040]
368[&!name="orchidaceae",!color=#008040]
375[&!name="tecophilaeaceae",!color=#008040]
376[&!name="doryanthaceae",!color=#008040]
377[&!name="iridaceae",!color=#008040]
379[&!name="xanthorrhoeaceae",!color=#008040]
380[&!name="amaryllidaceae",!color=#008040]
381[&!name="asparagaceae",!color=#008040]
382[&!name="arecaceae",!color=#008040]
384[&!name="commelinaceae",!color=#008040]
387[&!name="haemodoraceae",!color=#008040]
388[&!name="pontederiaceae",!color=#008040]
389[&!name="musaceae",!color=#008040]
390[&!name="heliconiaceae",!color=#008040]
392[&!name="lowiaceae",!color=#008040]
395[&!name="zingiberaceae",!color=#008040]
397[&!name="typhaceae",!color=#008040]
398[&!name="bromeliaceae",!color=#008040]
4[&!name="pinaceae",!color=#ff0000]
40[&!name="lauraceae",!color=#ff8000]
400[&!name="xyridaceae",!color=#008040]
410[&!name="poaceae",!color=#008040]
413[&!name="araceae",!color=#008040]
414[&!name="tofieldiaceae",!color=#008040]
415[&!name="alismataceae",!color=#008040]
416[&!name="hydrocharitaceae",!color=#008040]
417[&!name="butomaceae",!color=#008040]
419[&!name="aponogetonaceae",!color=#008040]
421[&!name="potamogetonaceae",!color=#008040]
425[&!name="juncaginaceae",!color=#008040]
43[&!name="papaveraceae",!color=#0080ff]
44[&!name="lardizabalaceae",!color=#0080ff]
47[&!name="berberidaceae",!color=#0080ff]
48[&!name="ranunculaceae",!color=#0080ff]
50[&!name="nelumbonaceae",!color=#66ffff]
51[&!name="platanaceae",!color=#66ffff]
53[&!name="trochodendraceae",!color=#0000ff]
54[&!name="buxaceae",!color=#0000ff]
56[&!name="gunneraceae",!color=#0000ff]
68[&!name="tamaricaceae",!color=#0000ff]
69[&!name="frankeniaceae",!color=#0000ff]
70[&!name="polygonaceae",!color=#0000ff]
71[&!name="plumbaginaceae",!color=#0000ff]
8[&!name="cupressaceae",!color=#ff0000]
81[&!name="amaranthaceae",!color=#0000ff]
83[&!name="caryophyllaceae",!color=#0000ff]
```

```
     88[&!name="aizoaceae",!color=#0000ff]
     9[&!name="taxaceae",!color=#ff0000]
     90[&!name="phytolaccaceae",!color=#0000ff]
     91[&!name="nyctaginaceae",!color=#0000ff]
     93[&!name="molluginaceae",!color=#0000ff]
     94[&!name="basellaceae",!color=#0000ff]
     98[&!name="talinaceae",!color=#0000ff]
     99[&!name="portulacaceae",!color=#0000ff]
;
end;

begin trees;
     tree tree_1 = [&R]
(((3[&!color=#ff0000]:270.6651,((4[&!color=#ff0000]:171.1527,(8[
&!color=#ff0000]:64.3894,9[&!color=#ff0000]:64.3894)[&!color=#ff
0000]:106.7633)[&!color=#ff0000]:55.7774,((10[&!color=#ff0000]:9
1.7523,11[&!color=#ff0000]:91.7523)[&!color=#ff0000]:55.333,12[&
!color=#ff0000]:147.0853)[&!color=#ff0000]:79.8448)[&!color=#ff0
000]:43.735)[&!color=#ff0000]:22.1938,2[&!color=#ff0000]:292.858
9)[&!color=#ff0000]:33.7966,(13[&!color=#400080,!rotate=false]:1
83.9897,(((17[&!color=#cc66ff,!rotate=false]:95.8563,19[&!color=
#cc66ff,!rotate=false]:95.8563)[&!color=#cc66ff,!rotate=false]:7
3.6687,((((43[&!color=#0080ff,!rotate=true]:109.78,(44[&!color=#
0080ff,!rotate=true]:86.2052,(47[&!color=#0080ff,!rotate=true]:5
5.7161,48[&!color=#0080ff,!rotate=true]:55.7161)[&!color=#0080ff
,!rotate=true]:30.4891)[&!color=#0080ff,!rotate=true]:23.5748)[&
!color=#0080ff,!rotate=true]:35.6683,((50[&!color=#66ffff,!rotat
e=true]:102.0917,51[&!color=#66ffff,!rotate=true]:102.0917)[&!co
lor=#66ffff,!rotate=true]:37.8791,(53[&!color=#0000ff,!rotate=tr
ue]:136.2739,(((((213[&!color=#000080,!rotate=true]:121.8314,(((
(214[&!color=#000080,!rotate=true]:96.7591,215[&!color=#000080,!
rotate=true]:96.7591)[&!color=#000080,!rotate=true]:15.0635,(217
[&!color=#000080,!rotate=true]:90.6845,((218[&!color=#000080,!ro
tate=true]:57.3663,219[&!color=#000080,!rotate=true]:57.3663)[&!
color=#000080,!rotate=true]:31.8412,(220[&!color=#000080,!rotate
=true]:83.2571,222[&!color=#000080,!rotate=true]:83.2561)[&!colo
r=#000080,!rotate=true]:5.9513)[&!color=#000080,!rotate=true]:1.
477)[&!color=#000080,!rotate=true]:21.1371)[&!color=#000080,!rot
ate=true]:2.9317,((229[&!color=#000080,!rotate=true]:27.4086,231
[&!color=#000080,!rotate=true]:27.4086)[&!color=#000080,!rotate=
true]:84.2929,(((239[&!color=#000080,!rotate=true]:61.7208,(240[
&!color=#000080,!rotate=true]:51.8619,(241[&!color=#000080,!rota
te=true]:46.2672,242[&!color=#000080,!rotate=true]:46.2672)[&!co
lor=#000080,!rotate=true]:5.5947)[&!color=#000080,!rotate=true]:
9.8589)[&!color=#000080,!rotate=true]:1.6186,237[&!color=#000080
,!rotate=true]:63.3395)[&!color=#000080,!rotate=true]:38.3513,((
257[&!color=#000080,!rotate=true]:79.4339,(261[&!color=#000080,!
rotate=true]:67.784,((268[&!color=#000080,!rotate=true]:46.0205,
```

(270[&!color=#000080,!rotate=true]:30.3737,(271[&!color=#000080,
!rotate=true]:22.1265,272[&!color=#000080,!rotate=true]:22.1265)
[&!color=#000080,!rotate=true]:8.2481)[&!color=#000080,!rotate=t
rue]:15.6469)[&!color=#000080,!rotate=true]:3.5921,267[&!color=#
000080,!rotate=true]:49.6136)[&!color=#000080,!rotate=true]:18.1
703)[&!color=#000080,!rotate=true]:11.6499)[&!color=#000080,!rot
ate=true]:11.0605,(247[&!color=#000080,!rotate=true]:67.3168,((2
48[&!color=#000080,!rotate=true]:56.5732,(249[&!color=#000080,!r
otate=true]:39.743,250[&!color=#000080,!rotate=true]:39.743)[&!c
olor=#000080,!rotate=true]:16.8302)[&!color=#000080,!rotate=true
]:6.77,(252[&!color=#000080,!rotate=true]:55.2428,253[&!color=#0
00080,!rotate=true]:55.2428)[&!color=#000080,!rotate=true]:8.100
4)[&!color=#000080,!rotate=true]:3.9737)[&!color=#000080,!rotate
=true]:23.1765)[&!color=#000080,!rotate=true]:11.1974)[&!color=#
000080,!rotate=true]:10.0108)[&!color=#000080,!rotate=true]:3.05
17)[&!color=#000080,!rotate=true]:2.5592,(((((((291[&!color=#000
080,!rotate=true]:52.842,293[&!color=#000080,!rotate=true]:52.84
1)[&!color=#000080,!rotate=true]:2.9788,289[&!color=#000080,!rot
ate=true]:55.8208)[&!color=#000080,!rotate=true]:42.153,(298[&!c
olor=#000080,!rotate=true]:58.0181,((299[&!color=#000080,!rotate
=true]:28.539,300[&!color=#000080,!rotate=true]:28.539)[&!color=
#000080,!rotate=true]:13.4866,303[&!color=#000080,!rotate=true]:
42.0256)[&!color=#000080,!rotate=true]:15.9935)[&!color=#000080,
!rotate=true]:39.9556)[&!color=#000080,!rotate=true]:3.5135,(280
[&!color=#000080,!rotate=true]:83.2662,((285[&!color=#000080,!ro
tate=true]:56.5445,(286[&!color=#000080,!rotate=true]:44.2088,(2
87[&!color=#000080,!rotate=true]:34.5194,288[&!color=#000080,!ro
tate=true]:34.5194)[&!color=#000080,!rotate=true]:9.6894)[&!colo
r=#000080,!rotate=true]:12.3357)[&!color=#000080,!rotate=true]:1
1.9465,283[&!color=#000080,!rotate=true]:68.49)[&!color=#000080,
!rotate=true]:14.7762)[&!color=#000080,!rotate=true]:18.22)[&!co
lor=#000080,!rotate=true]:1.9956,(276[&!color=#000080,!rotate=tr
ue]:59.3632,278[&!color=#000080,!rotate=true]:59.3632)[&!color=#
000080,!rotate=true]:44.1196)[&!color=#000080,!rotate=true]:8.11
79,(((((327[&!color=#000080,!rotate=true]:86.2324,(340[&!color=#
000080,!rotate=true]:72.3138,(342[&!color=#000080,!rotate=true]:
59.3003,((343[&!color=#000080,!rotate=true]:46.7128,344[&!color=
#000080,!rotate=true]:46.7128)[&!color=#000080,!rotate=true]:10.
9498,347[&!color=#000080,!rotate=true]:57.6626)[&!color=#000080,
!rotate=true]:1.6377)[&!color=#000080,!rotate=true]:13.0135)[&!c
olor=#000080,!rotate=true]:13.9185)[&label="B",!color=#000080,!r
otate=true]:11.4435,325[&!color=#000080,!rotate=true]:97.6758)[&
!color=#000080,!rotate=true]:2.8372,((318[&!color=#000080,!rotat
e=true]:94.9126,(321[&!color=#000080,!rotate=true]:87.0564,322[&
!color=#000080,!rotate=true]:87.0564)[&!color=#000080,!rotate=tr
ue]:7.8562)[&!color=#000080,!rotate=true]:2.96,316[&!color=#0000
80,!rotate=true]:97.8726)[&!color=#000080,!rotate=true]:2.6414)[
&!color=#000080,!rotate=true]:5.8939,((307[&!color=#000080,!rota

te=true]:36.8778,308[&!color=#000080,!rotate=true]:36.8778)[&!co
lor=#000080,!rotate=true]:24.7964,(309[&!color=#000080,!rotate=t
rue]:47.4203,310[&!color=#000080,!rotate=true]:47.4203)[&!color=
#000080,!rotate=true]:14.2539)[&!color=#000080,!rotate=true]:44.
7346)[&!color=#000080,!rotate=true]:1.077,305[&!color=#000080,!r
otate=true]:107.4848)[&!color=#000080,!rotate=true]:4.1159)[&!co
lor=#000080,!rotate=true]:1.858,273[&!color=#000080,!rotate=true
]:113.4587)[&!color=#000080,!rotate=true]:3.8537)[&!color=#00008
0,!rotate=true]:4.52)[&!color=#000080,!rotate=true]:1.7341,((200
[&!color=#0000ff,!rotate=true]:80.5335,(202[&!color=#0000ff,!rot
ate=true]:60.0028,203[&!color=#0000ff,!rotate=true]:60.0018)[&!c
olor=#0000ff,!rotate=true]:20.5307)[&!color=#0000ff,!rotate=true
]:3.2827,((208[&!color=#0000ff,!rotate=true]:64.6079,(211[&!colo
r=#0000ff,!rotate=true]:30.4605,212[&!color=#0000ff,!rotate=true
]:30.4605)[&!color=#0000ff,!rotate=true]:34.1464)[&!color=#0000f
f,!rotate=true]:15.2555,(206[&!color=#0000ff,!rotate=true]:43.89
22,207[&!color=#0000ff,!rotate=true]:43.8922)[&!color=#0000ff,!r
otate=true]:35.9702)[&!color=#0000ff,!rotate=true]:3.9528)[&!col
or=#0000ff,!rotate=true]:39.7493)[&!color=#0000ff,!rotate=true]:
2.2604,((((68[&!color=#0000ff,!rotate=true]:43.4508,69[&!color=#
0000ff,!rotate=true]:43.4508)[&!color=#0000ff,!rotate=true]:36.4
343,(70[&!color=#0000ff,!rotate=true]:48.7324,71[&!color=#0000ff
,!rotate=true]:48.7324)[&!color=#0000ff,!rotate=true]:31.1527)[&
!color=#0000ff,!rotate=true]:14.6146,((81[&!color=#0000ff,!rotat
e=true]:53.1362,83[&!color=#0000ff,!rotate=true]:53.1372)[&!colo
r=#0000ff,!rotate=true]:13.344,((93[&!color=#0000ff,!rotate=true
]:47.1586,(94[&!color=#0000ff,!rotate=true]:36.015,(98[&!color=#
0000ff,!rotate=true]:25.1098,(99[&!color=#0000ff,!rotate=true]:2
1.1636,101[&!color=#0000ff,!rotate=true]:21.1636)[&!color=#0000f
f,!rotate=true]:3.9462)[&!color=#0000ff,!rotate=true]:10.9053)[&
!color=#0000ff,!rotate=true]:11.1436)[&!color=#0000ff,!rotate=tr
ue]:7.765,(88[&!color=#0000ff,!rotate=true]:35.282,(90[&!color=#
0000ff,!rotate=true]:28.2974,91[&!color=#0000ff,!rotate=true]:28
.2974)[&!color=#0000ff,!rotate=true]:6.9847)[&!color=#0000ff,!ro
tate=true]:19.6415)[&!color=#0000ff,!rotate=true]:11.5567)[&!col
or=#0000ff,!rotate=true]:28.0204)[&!color=#0000ff,!rotate=true]:
19.9598,((((111[&!color=#0000ff,!rotate=true]:69.4548,((112[&!co
lor=#0000ff,!rotate=true]:54.0609,113[&!color=#0000ff,!rotate=tr
ue]:54.0609)[&!color=#0000ff,!rotate=true]:12.2736,((114[&!color
=#0000ff,!rotate=true]:59.141,(115[&!color=#0000ff,!rotate=true]
:53.0118,116[&!color=#0000ff,!rotate=true]:53.0118)[&!color=#000
0ff,!rotate=true]:6.1292)[&!color=#0000ff,!rotate=true]:3.8985,(
120[&!color=#0000ff,!rotate=true]:56.6167,((122[&!color=#0000ff,
!rotate=true]:39.3478,(123[&!color=#0000ff,!rotate=true]:30.8951
,124[&!color=#0000ff,!rotate=true]:30.8951)[&!color=#0000ff,!rot
ate=true]:8.4527)[&!color=#0000ff,!rotate=true]:7.2371,(125[&!co
lor=#0000ff,!rotate=true]:38.7437,127[&!color=#0000ff,!rotate=tr
ue]:38.7437)[&!color=#0000ff,!rotate=true]:7.8412)[&!color=#0000

ff,!rotate=true]:10.0318)[&!color=#0000ff,!rotate=true]:6.4227)[
&!color=#0000ff,!rotate=true]:3.2941)[&!color=#0000ff,!rotate=tr
ue]:3.1213)[&!color=#0000ff,!rotate=true]:23.515,108[&!color=#00
00ff,!rotate=true]:92.9698)[&!color=#0000ff,!rotate=true]:11.289
1,((((((138[&!color=#0000ff,!rotate=true]:40.3522,(140[&!color=#
0000ff,!rotate=true]:18.7465,141[&!color=#0000ff,!rotate=true]:1
8.7465)[&!color=#0000ff,!rotate=true]:21.6056)[&!color=#0000ff,!
rotate=true]:8.9609,142[&!color=#0000ff,!rotate=true]:49.3131)[&
!color=#0000ff,!rotate=true]:9.457,137[&!color=#0000ff,!rotate=t
rue]:58.7701)[&!color=#0000ff,!rotate=true]:17.2638,((150[&!colo
r=#0000ff,!rotate=true]:41.4061,151[&!color=#0000ff,!rotate=true
]:41.4061)[&!color=#0000ff,!rotate=true]:21.0377,(155[&!color=#0
000ff,!rotate=true]:29.4913,(156[&!color=#0000ff,!rotate=true]:2
6.7144,158[&!color=#0000ff,!rotate=true]:26.7144)[&!color=#0000f
f,!rotate=true]:2.7768)[&!color=#0000ff,!rotate=true]:32.9535)[&
!color=#0000ff,!rotate=true]:13.5892)[&!color=#0000ff,!rotate=tr
ue]:16.4615,134[&!color=#0000ff,!rotate=true]:92.4954)[&!color=#
0000ff,!rotate=true]:5.0256,(161[&!color=#0000ff,!rotate=true]:8
9.5513,((165[&!color=#0000ff,!rotate=true]:56.0489,(166[&!color=
#0000ff,!rotate=true]:43.5349,(168[&!color=#0000ff,!rotate=true]
:34.0727,169[&!color=#0000ff,!rotate=true]:34.0727)[&!color=#000
0ff,!rotate=true]:9.4622)[&!color=#0000ff,!rotate=true]:12.514)[
&!color=#0000ff,!rotate=true]:15.5422,(170[&!color=#0000ff,!rota
te=true]:68.0705,((171[&!color=#0000ff,!rotate=true]:37.4597,172
[&!color=#0000ff,!rotate=true]:37.4597)[&!color=#0000ff,!rotate=
true]:29.0688,(177[&!color=#0000ff,!rotate=true]:50.9547,((182[&
!color=#0000ff,!rotate=true]:38.6666,(183[&!color=#0000ff,!rotat
e=true]:36.0308,(184[&!color=#0000ff,!rotate=true]:34.4251,((185
[&!color=#0000ff,!rotate=true]:29.4367,(186[&!color=#0000ff,!rot
ate=true]:26.8774,(187[&!color=#0000ff,!rotate=true]:22.112,188[
&!color=#0000ff,!rotate=true]:22.112)[&!color=#0000ff,!rotate=tr
ue]:4.7654)[&!color=#0000ff,!rotate=true]:2.5593)[&!color=#0000f
f,!rotate=true]:3.1899,((((196[&!color=#0000ff,!rotate=true]:25.
9174,198[&!color=#0000ff,!rotate=true]:25.9174)[&!color=#0000ff,
!rotate=true]:2.5591,193[&!color=#0000ff,!rotate=true]:28.4765)[
&!color=#0000ff,!rotate=true]:1.1342,191[&!color=#0000ff,!rotate
=true]:29.6107)[&!color=#0000ff,!rotate=true]:1.079,190[&!color=
#0000ff,!rotate=true]:30.6896)[&!color=#0000ff,!rotate=true]:1.9
37)[&!color=#0000ff,!rotate=true]:1.7985)[&!color=#0000ff,!rotat
e=true]:1.6057)[&!color=#0000ff,!rotate=true]:2.6358)[&!color=#0
000ff,!rotate=true]:1.3669,181[&!color=#0000ff,!rotate=true]:40.
0325)[&!color=#0000ff,!rotate=true]:10.9212)[&!color=#0000ff,!ro
tate=true]:15.5748)[&!color=#0000ff,!rotate=true]:1.541)[&!color
=#0000ff,!rotate=true]:3.5216)[&!color=#0000ff,!rotate=true]:17.
9592)[&!color=#0000ff,!rotate=true]:7.9707)[&!color=#0000ff,!rot
ate=true]:6.7379)[&!color=#0000ff,!rotate=true]:1.7852,(103[&!co
lor=#0000ff,!rotate=true]:69.2055,(104[&!color=#0000ff,!rotate=t
rue]:38.3012,105[&!color=#0000ff,!rotate=true]:38.3012)[&!color=

#0000ff,!rotate=true]:30.9033)[&!color=#0000ff,!rotate=true]:36.
8386)[&!color=#0000ff,!rotate=true]:8.4153)[&!color=#0000ff,!rot
ate=true]:11.3665)[&!color=#0000ff,!rotate=true]:2.7873,56[&!col
or=#0000ff,!rotate=true]:128.6132)[&!color=#0000ff,!rotate=true]
:5.5109,54[&!color=#0000ff,!rotate=true]:134.1241)[&!color=#0000
ff,!rotate=true]:2.1498)[&!color=#0000ff,!rotate=true]:3.6969)[&
!color=#0000ff,!rotate=true]:5.4775)[&!color=#0000ff,!rotate=tru
e]:14.2652,((413[&!color=#008040,!rotate=true]:121.5893,(414[&!c
olor=#008040,!rotate=true]:116.0839,((415[&!color=#008040,!rotat
e=true]:72.877,(416[&!color=#008040,!rotate=true]:65.5031,417[&!
color=#008040,!rotate=true]:65.5031)[&!color=#008040,!rotate=tru
e]:7.3739)[&!color=#008040,!rotate=true]:21.7587,(419[&!color=#0
08040,!rotate=true]:81.4526,(425[&!color=#008040,!rotate=true]:6
3.1043,421[&!color=#008040,!rotate=true]:63.1043)[&!color=#00804
0,!rotate=true]:18.3483)[&!color=#008040,!rotate=true]:13.182)[&
!color=#008040,!rotate=true]:21.4483)[&!color=#008040,!rotate=tr
ue]:5.5054)[&!color=#008040,!rotate=true]:13.9164,((352[&!color=
#008040,!rotate=true]:92.9267,(354[&!color=#008040,!rotate=true]
:57.9442,357[&!color=#008040,!rotate=true]:57.9452)[&!color=#008
040,!rotate=true]:34.9825)[&!color=#008040,!rotate=true]:25.2033
,(((368[&!color=#008040,!rotate=true]:99.0365,((376[&!color=#008
040,!rotate=true]:62.2398,(377[&!color=#008040,!rotate=true]:52.
5609,(379[&!color=#008040,!rotate=true]:34.6323,(380[&!color=#00
8040,!rotate=true]:28.5824,381[&!color=#008040,!rotate=true]:28.
5824)[&!color=#008040,!rotate=true]:6.0499)[&!color=#008040,!rot
ate=true]:17.9287)[&!color=#008040,!rotate=true]:9.6788)[&!color
=#008040,!rotate=true]:3.0008,375[&!color=#008040,!rotate=true]:
65.2416)[&!color=#008040,!rotate=true]:33.7949)[&!color=#008040,
!rotate=true]:12.9637,((382[&!color=#008040,!rotate=true]:104.21
24,((384[&!color=#008040,!rotate=true]:76.1336,(387[&!color=#008
040,!rotate=true]:55.6399,388[&!color=#008040,!rotate=true]:55.6
399)[&!color=#008040,!rotate=true]:20.4938)[&!color=#008040,!rot
ate=true]:17.6002,(389[&!color=#008040,!rotate=true]:84.6442,((3
90[&!color=#008040,!rotate=true]:61.4558,392[&!color=#008040,!ro
tate=true]:61.4558)[&!color=#008040,!rotate=true]:12.7915,395[&!
color=#008040,!rotate=true]:74.2473)[&!color=#008040,!rotate=tru
e]:10.3969)[&!color=#008040,!rotate=true]:9.0887)[&!color=#00804
0,!rotate=true]:10.4796)[&!color=#008040,!rotate=true]:2.453,((3
97[&!color=#008040,!rotate=true]:79.4529,398[&!color=#008040,!ro
tate=true]:79.4529)[&!color=#008040,!rotate=true]:10.829,(400[&!
color=#008040,!rotate=true]:81.8867,410[&!color=#008040,!rotate=
true]:81.8877)[&!color=#008040,!rotate=true]:8.3951)[&!color=#00
8040,!rotate=true]:16.3826)[&!color=#008040,!rotate=true]:5.3348
)[&!color=#008040,!rotate=true]:3.5025,((358[&!color=#008040,!ro
tate=true]:81.4059,(360[&!color=#008040,!rotate=true]:51.561,361
[&!color=#008040,!rotate=true]:51.561)[&!color=#008040,!rotate=t
rue]:29.8449)[&!color=#008040,!rotate=true]:5.6129,(364[&!color=
#008040,!rotate=true]:40.9678,365[&!color=#008040,!rotate=true]:

```
40.9678)[&!color=#008040,!rotate=true]:46.05)[&!color=#008040,!r
otate=true]:28.4839)[&!color=#008040,!rotate=true]:2.6273)[&!col
or=#008040,!rotate=true]:17.3756)[&!color=#008040,!rotate=true]:
24.2089)[&!rotate=true]:1.059,(((34[&!color=#ff8000,!rotate=true
]:110.6748,40[&!color=#ff8000,!rotate=true]:110.6748)[&!color=#f
f8000,!rotate=true]:13.5402,(29[&!color=#ff8000,!rotate=true]:58
.2596,32[&!color=#ff8000,!rotate=true]:58.2596)[&!color=#ff8000,
!rotate=true]:65.9554)[&!color=#ff8000,!rotate=true]:9.4618,((26
[&!color=#ff8000,!rotate=true]:54.2685,27[&!color=#ff8000,!rotat
e=true]:54.2685)[&!color=#ff8000,!rotate=true]:48.4033,24[&!colo
r=#ff8000,!rotate=true]:102.6718)[&!color=#ff8000,!rotate=true]:
31.005)[&!color=#ff8000,!rotate=true]:27.0967)[&!rotate=true]:8.
7515)[&!rotate=false]:8.6069,(15[&!color=#ff00ff,!rotate=true]:3
2.54,16[&!color=#ff00ff,!rotate=true]:32.54)[&!color=#ff00ff,!ro
tate=true]:145.5919)[&!rotate=true]:5.8577)[&!rotate=false]:142.
6659);
end;

begin figtree;
     set appearance.backgroundColorAttribute="Default";
     set appearance.backgroundColour=#ffffff;
     set appearance.branchColorAttribute="User selection";
     set appearance.branchColorGradient=false;
     set appearance.branchLineWidth=1.0;
     set appearance.branchMinLineWidth=0.0;
     set appearance.branchWidthAttribute="Fixed";
     set appearance.foregroundColour=#000000;
     set appearance.hilightingGradient=false;
     set appearance.selectionColour=#2d3680;
     set branchLabels.colorAttribute="User selection";
     set branchLabels.displayAttribute="Branch times";
     set branchLabels.fontName="sansserif";
     set branchLabels.fontSize=8;
     set branchLabels.fontStyle=0;
     set branchLabels.isShown=false;
     set branchLabels.significantDigits=4;
     set layout.expansion=0;
     set layout.layoutType="RECTILINEAR";
     set layout.zoom=0;
     set legend.attribute="label";
     set legend.fontSize=10.0;
     set legend.isShown=false;
     set legend.significantDigits=4;
     set nodeBars.barWidth=4.0;
     set nodeBars.displayAttribute=null;
     set nodeBars.isShown=false;
     set nodeLabels.colorAttribute="User selection";
     set nodeLabels.displayAttribute="Node ages";
```

```
set nodeLabels.fontName="sansserif";
set nodeLabels.fontSize=8;
set nodeLabels.fontStyle=0;
set nodeLabels.isShown=false;
set nodeLabels.significantDigits=4;
set nodeShapeExternal.colourAttribute="User selection";
set nodeShapeExternal.isShown=false;
set nodeShapeExternal.minSize=10.0;
set nodeShapeExternal.scaleType=Width;
set nodeShapeExternal.shapeType=Circle;
set nodeShapeExternal.size=4.0;
set nodeShapeExternal.sizeAttribute="Fixed";
set nodeShapeInternal.colourAttribute="User selection";
set nodeShapeInternal.isShown=false;
set nodeShapeInternal.minSize=10.0;
set nodeShapeInternal.scaleType=Width;
set nodeShapeInternal.shapeType=Circle;
set nodeShapeInternal.size=4.0;
set nodeShapeInternal.sizeAttribute="Fixed";
set polarLayout.alignTipLabels=false;
set polarLayout.angularRange=0;
set polarLayout.rootAngle=0;
set polarLayout.rootLength=100;
set polarLayout.showRoot=true;
set radialLayout.spread=0.0;
set rectilinearLayout.alignTipLabels=false;
set rectilinearLayout.curvature=0;
set rectilinearLayout.rootLength=100;
set scale.offsetAge=0.0;
set scale.rootAge=1.0;
set scale.scaleFactor=1.0;
set scale.scaleRoot=false;
set scaleAxis.automaticScale=true;
set scaleAxis.fontSize=8.0;
set scaleAxis.isShown=false;
set scaleAxis.lineWidth=1.0;
set scaleAxis.majorTicks=1.0;
set scaleAxis.minorTicks=0.5;
set scaleAxis.origin=0.0;
set scaleAxis.reverseAxis=false;
set scaleAxis.showGrid=true;
set scaleBar.automaticScale=true;
set scaleBar.fontSize=10.0;
set scaleBar.isShown=true;
set scaleBar.lineWidth=1.0;
set scaleBar.scaleRange=0.0;
set tipLabels.colorAttribute="User selection";
set tipLabels.displayAttribute="Names";
```

```
    set tipLabels.fontName="sansserif";
    set tipLabels.fontSize=7;
    set tipLabels.fontStyle=0;
    set tipLabels.isShown=true;
    set tipLabels.significantDigits=4;
    set trees.order=false;
    set trees.orderType="increasing";
    set trees.rooting=false;
    set trees.rootingType="User Selection";
    set trees.transform=false;
    set trees.transformType="cladogram";
end;
```
